# Glycosylation of Zika Virus Is Important in Host-Virus Interaction and Pathogenesis

**DOI:** 10.1101/564542

**Authors:** Nanda Kishore Routhu, Sylvain D. Lehoux, Emily A. Rouse, Mehdi R. M. Bidokhti, Leila B. Giron, Alit Anzurez, St Patrick Reid, Mohamed Abdel-Mohsen, Richard D. Cummings, Siddappa N. Byrareddy

## Abstract

Zika virus (ZIKV) is a global public health issue due to its association with severe developmental disorders in infants and neurological disorders in adults. Because ZIKV uses glycosylation of its envelope (E) protein to interact with host cell receptors to facilitate entry, these interactions could also be important for designing therapeutics and vaccines. Due to a lack of information about Asn-linked (N-glycans) on ZIKV E, we analyzed ZIKV E of various strains derived from different cells. ZIKV E proteins are extensively modified with oligomannose-, hybrid- and complex-N-glycans of a highly heterogeneous nature. Host cell-surface glycans correlated strongly with the glycomic features of ZIKV E. Mechanistically, we discovered that ZIKV N-glycans are important in viral pathogenesis, as mannose-specific C-type lectins DC-SIGN and L-SIGN mediate cell entry of ZIKV. Our findings represent the first detailed mapping of N-glycans on ZIKV E of various strains and their functional significance.

## Introduction

Zika virus (ZIKV) is mainly transmitted to humans *via* infected mosquitoes, though other transmission routes, such as through placenta and sexual intercourse can also occur ^1,2^ Recently, the considerable increase in ZIKV infection rates has raised urgent global urgent concerns, especially after the outbreaks in Yap Islands (2007) ^3,4^ French Polynesia (2013) ^5,6^, Easter Island (2014) ^7^, and Brazil (2015) ^8,9^ These ZIKV outbreaks were associated with a sharp increase in cases of Guillain Barre Syndrome (GBS), an autoimmune disease featured by weakening and paralysis of the limbs and face. In 2015, Zika spread to South and Central America, infecting thousands of people in Brazil and Colombia, where it was associated with an increase in GBS rates as well as a significant increase in severe fetal abnormalities including spontaneous abortion, stillbirth, hydrocephaly, microcephaly, hydranencephaly/hydrops fetalis, and placental insufficiency ^10–14^ ZIKV is a small, enveloped positive-strand RNA virus ^15^. The genomic RNA of this mosquito-borne flavivirus contains a single open reading frame (ORF) that encodes a polyprotein ^16,17^ This polyprotein undergoes co- and post-translational processing to produce three structural proteins, which are capsid (C), pre-membrane (prM) and envelope (E), in addition to seven non-structural proteins (NS1, NS2A, NS2B, NS3, NS4A, NS4B, and NS5) ^17,18^. Of the three structural proteins, E is the major surface glycoprotein, containing three domains (I, II, III) and two transmembrane helices ^17–19^. The majority of flaviviruses E proteins are post-translationally modified by N-glycans at amino acid 153/154 within a highly conserved glycosylation motif of N-X-T/S at positions 154-156 (where X is any amino acid except proline) ^16^. This is a primary target of neutralizing antibodies and is required for virus entry ^20^.

Previously, it was demonstrated that the N-glycosylation sites on the E protein of flaviviruses are highly conserved **(Supplementary Fig. 1)**, playing a vital role in both infectivity and assembly ^16,21–23^. Loss of N-glycosylation motif on domain I of E protein results in impaired expression and secretion of E ectodomain from mammalian cells ^24^. Furthermore, studies in mice have shown that ZIKVs lacking the N-glycans of the ZIKV E were severely compromised for their ability to cause mortality and neuroinvasion, suggesting a vital role for these glycans of the E protein in ZIKV pathogenesis ^22,25^. In this study, we utilized mass spectrometry (MS) and a lectin microarray to analyze the structures and composition of glycans on ZIKV E from different strains and on the surface of the virus-producing cells, as well as to explore mosquito-human mode of ZIKV transmission and its association to neurological disorders. Importantly, our functional studies show that these N-glycans of the ZIKV envelope glycoprotein are important in viral infection of cells. Our findings have implications on understanding the roles for these glycan patterns on ZIKV E in virus-host interaction and pathogenesis.

## Results

### Glycoprotein E localizes subcellularly in ZIKV-infected cells

In order to confirm whether ZIKV E localized within the endoplasmic reticulum (ER), mammalian cells (Vero and SNB19 cells) and an insect cell line (C6/36 cells) were infected with the following ZIKV strains 1) Asian strains (PRVABC59 and FLR isolates), 2) African strains (MR766 and IbH isolates) and 3) a Brazilian isolate from 2016 (SJRP) for 24 hours. As shown in **Fig. 1**, the patterns of fluorescence demonstrated that ZIKV E localizes to the periphery of cell nucleus, consistent with localization in the ER. The cytoplasmic area of Vero cells and SNB19 cells that were infected with PRVABC59 and MR766 displayed similar patterns of perinuclear and nuclear immunofluorescence of ZIKV E. However, C6/36 cells are smaller in size compared to other cell lines and displayed reduced cytoplasmic area compared to uninfected cells. The cells that were infected with either the MR766 or IbH strain exhibited particularly reduced cytoplasmic area. These data suggest that the ZIKV E localizes predominantly to ER.

**Fig. 1.**
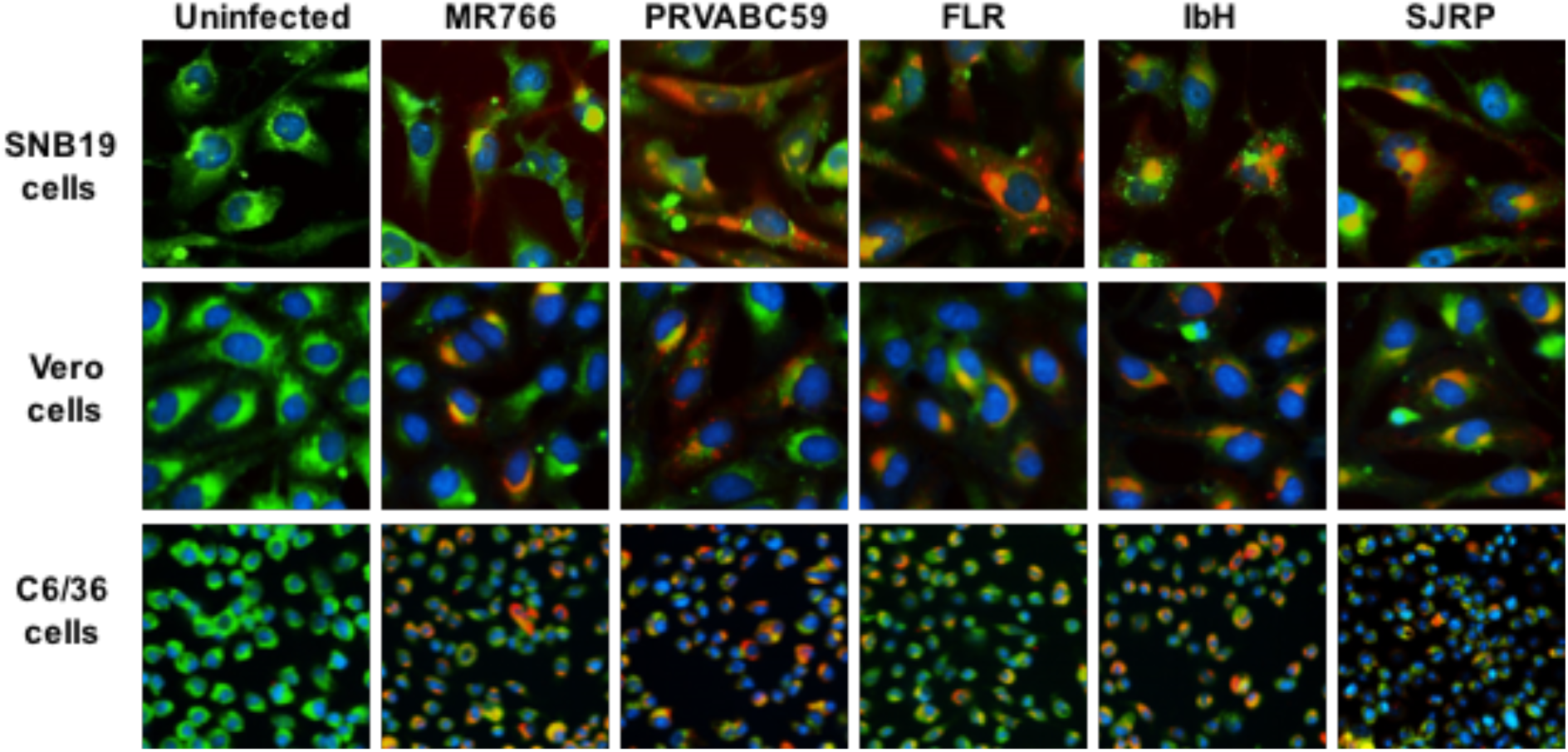
Analyzing the ER-localization of the ZIKV E glycoprotein in infected cells. Immunofluorescence images of SNB-19 cells, Vero cells, and C6/36 cells that were infected with 0.1 MOI of ZIKV (MR766, PRVABC59, FLR, IbH, and SJRP). After 24 hours, the cells were fixed, permeabilized, and stained for E protein using a monoclonal anti-flavivirus group antigen antibody (clone D1-4G2-4-15) (red), ER using ER ID dye (green), and cell nuclei using Hoechst stain (blue). No red fluorescence is observed in control cells.

### MS analysis of ZIKV E derived from different cell lines showed various profiles of N-glycan pattern

Following the production and purification of the different ZIKV strains in various cell lines, the virions were pelleted, lysed, resolved in 12% bis-tris protein gels, and stained with Coomassie Blue. The bands corresponding to the ZIKV E were excised for N-glycan MS profiling **(Supplementary Fig. 3)**. The relative abundances of the N-glycans released by PNGase F, and identified for each of the five ZIKV E proteins produced in each of the six different cell lines, are presented as a heatmap in **Fig. 2**; the predicted glycan structures with the corresponding sizes are presented in **Supplementary Table 1**. Although distinct N-glycosylation patterns emerged, the two ZIKV E proteins from African lineage strains, MR766 and IbH, displayed similar N-glycan profiles across the six cell lines that were tested **(Supplementary Fig. 4a, b)**. When produced in C6/36, Vero, THP-1, and SNB19 cells, these N-glycan profiles were heavily dominated by sialylated glycans. For these four cell lines, bi- and tri-sialylated tri-antennary N-glycans (3241.8 m/z and 3603.0 m/z) were the most abundant found on ZIKV E, followed by bi-sialylated bi-antennary (2792.5 m/z) and tetra-sialylated tri-antennary (3964.2 m/z) N-glycans **(Supplementary Fig. 4a, b)**. ZIKV E produced in SNB19 cells was also found to be decorated with Man5 (1579.9 m/z), agalactosylated bisected (1907.1 m/z), core-fucosylated non-sialylated bi-antennary (2244.3 m/z), and core-fucosylated mono-sialylated bi-antennary (2605.5 m/z) N-glycans **(Supplementary Fig. 4a, b)**. When expressed in the LLC-MK2 and JEG-9 cell lines, the N-glycan profiles of MR766 and IbH strains of E were markedly different. When produced in LLC-MK2 cells, bi-sialylated bi-antennary (2792.5 m/z) and core-fucosylated agalactosylated (1836.0 m/z) N-glycans were by far the two most abundant glycan species **(Supplementary Fig. 4a, b)**. More complex N-glycans, with three or more sialic acids and additional lactosamine repeats, were not identified in ZIKV E expressed in LLC-MK2 cells. Finally, the ZIKV E produced in JEG-3 cells revealed a different N-glycosylation pattern between MR766 and IbH strains. While the di- and tri-sialylated N-glycans (2792.5, 3241.8, and 3603.0 m/z) were the most abundant for the MR766 ZIKV-E produced in JEG-3, the variety of N-glycans identified in JEG-3 cells was greater, notably including Man_5_GlcNAc_2_ (1579.9 m/z), Man_6_GlcNAc_2_ (1784.0 m/z), Man_7_GlcNAc_2_ (1988.1 m/z), Man_8_GlcNAc_2_ (2192.2 m/z), asialo bi-antennary (2070.2 m/z), and core fucosylated asialo-bi-antennary (2244.3 m/z) N-glycans **(Supplementary Fig. 4a)**. In contrast, the IbH strain produced in the JEG-3 cells displayed a more restricted N-glycan profile, with the notable absence of sialylated N-glycans and with the core fucosylated asialo-bi-antennary (2244.3 m/z) N-glycan being the most complex N-glycan identified (**Supplementary Fig. 4b**).

**Fig. 2.**
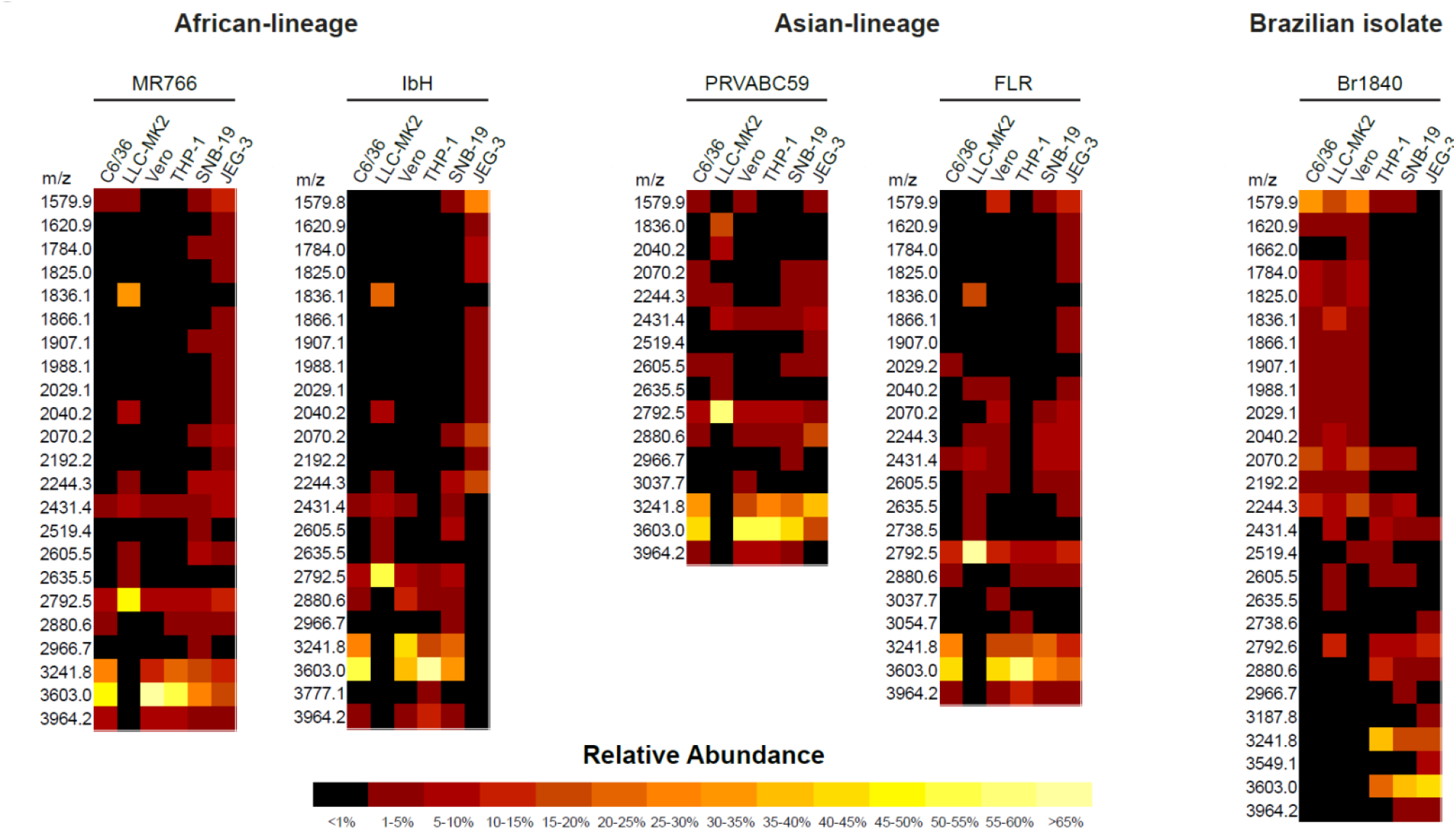
Heatmap presentation of N-glycans of glycoprotein E of purified ZIKV. ZIKVs (MR766, IbH, PRVABC59, FLR, and SJRP) produced from various cell types (C6/36, LLC-MK2, Vero, THP-1, SNB-19, and JEG-3 cells) were purified using sucrose cushion ultracentrifugation and resolved on SDS-PAGE. The presence of glycoprotein E was confirmed through western blotting. Glycoprotein E was then excised and subjected to N-glycan release, followed by MS identification. The N-glycans identified through this method are represented in heatmap format.

Similar to African lineage ZIKV, the N-glycosylation profiles from the Asian lineage viruses PRVABC59 and FLR varied across cell lines, yet were nearly identical within any given cell line (**Supplementary Fig. 4c, d**). Additionally, the E protein of PRVABC59 and FLR strains expressed in C6/36, Vero, THP-1, and SNB19 cell lines were primarily decorated with tri-sialylated tri-antennary N-glycans (3603.0 m/z) and—with a lesser abundance—di-sialylated bi- and tri-antennary (2792.5 and 3241.8 m/z) and tetra-sialylated tri-antennary (3964.2 m/z) N-glycans (**Supplementary Fig. 4c**, d). Minor N-glycans including Man_5_GlcNAc_2_ (1579.9 m/z), monosialylated bi-antennary (2431.4 m/z), core-fucosylated mono-galactosylated bi-antennary (2040.2 m/z), core-fucosylated asialo bi-antennary (2244.3 m/z), and core-fucosylated monosialylated bi-antennary (2605.5 m/z) N-glycans were also identified (**Supplementary Fig. 4c, d**). Similar to the African lineage, the E proteins of Asian lineage revealed a more restricted N-glycan profile when expressed in LLC-MK2 cells. The di-sialylated bi-antennary (2792.5 m/z) N-glycan was identified as the most abundant N-glycan, with very low to no expression of more complex N-glycans. The core-fucosylated agalactosylated (1836.0 m/z) and mono-sialylated bi-antennary (2431.4 m/z) were, respectively, the second and third most relatively abundant N-glycans reported (**Supplementary Fig. 4c, d**). Lastly, the N-glycan profiles of the PRVABC59 and FLR strains of E expressed in JEG-3 cells showed a lower abundance of highly sialylated N-glycans such as the tri- and tetra-sialylated tri-antennary N-glycans (3603.0 and 3964.2 m/z) compared to the virus produced in the C6/36, Vero, THP-1, and SNB19 cells. By contrast, the E of the PRVABC59 strain showed an increased abundance of both mono-sialylated N-glycans (2431.4 and 2880.6 m/z) and non-sialylated N-glycans (2040.2, 2070.2 and 2244.3 m/z).

Unlike the African and Asian lineage strains, the N-glycans found on the E protein of the Brazilian isolate SJRP were almost exclusively non-sialylated when produced in C6/36 and Vero cells. In both of these cell lines, the major N-glycans found were Man_5_GlcNAc_2_ (1579.9 m/z), asialo-bi-antennary (2070.2 m/z), and core fucosylated asialo-bi-antennary (2244.3 m/z) N-glycans (**Supplementary Fig. 4e**). When produced in LLC-MK2 cells, however, the N-glycan profile of the E protein from the SJRP strain was comparable to the profiles found for the four other strains, with the di-sialylated bi-antennary (2792.6 m/z) and the core-fucosylated agalactosylated (1836.1 m/z) N-glycans being the most abundant. Contrasting the other E proteins produced in C6/36, LLC-MK2 and Vero cells, Man5 (1579.9 m/z) was found to be the most abundant N-glycan on the SJRP strain. However, the N-glycan profiles generated from the E protein of SJRP strain expressed in THP-1, SNB19, and JEG-3 cells revealed a high relative abundance of di-sialylated bi-antennary (2792.5 m/z), tri-antennary (3241.8 m/z), and tri-sialylated tri-antennary (3603.0 m/z) with little to no expression of less complex non-sialylated N-glycans such as Man5GlcNAc2 (**Supplementary Fig. 4e**).

The major proportion (> 5%) of N-glycan spectra at 3603.0 mz, 3241.8 mz, 2792.6 mz, 3964.2 mz, 1836.0 mz, and 2040.2 mz were identified on ZIKV E produced in these cell lines. These heterogenous N-glycan structures of the studied ZIKV E include high (oligo)-mannose; these and other N-glycans compositionally are predicted to include the residues N-acetylgalactosamine (GalNAc), N-acetylglucosamine (GlcNAc), N-acetylneuraminic acid (NeuAc), galactose (Gal), glucose (Glc), fucose (Fuc), and sialic acid. In addition, we found many other glycan forms with high heterogeneity but smaller proportion (< 5%); these were not further characterized. These data are indicative of high heterogeneity of structure and composition of N-glycans on ZIKV E proteins (**Supplementary Table 2**).

### Cell-surface glycosylation strongly correlates with glycomic features of ZIKV E protein

In order to identify the general patterns of endogenous glycans in ZIKV-producing cell surface endogenous glycoproteins, a lectin array was used. This approach relies on specificities of immobilized lectins that recognize specific glycan features on cell surface glycoproteins, resulting in their binding to the array. These data revealed that different cell lines harbor differential cell-surface glycosylation **(Fig. 3)** that might impact the glycosylation of the virus produced in these cells. In general, the expression of several cell-surface glycan structures correlated with viral N-glycan features (**Supplementary Table 3**). Of interest, and as shown in **Fig. 4**, strong correlation between the presence of cell-surface Fuc, sialic acid, lactose, mannose, GlcNAc, and the N-glycan features of E-protein. There was no statistically significant difference with other proteins analyzed.

**Fig. 3.**
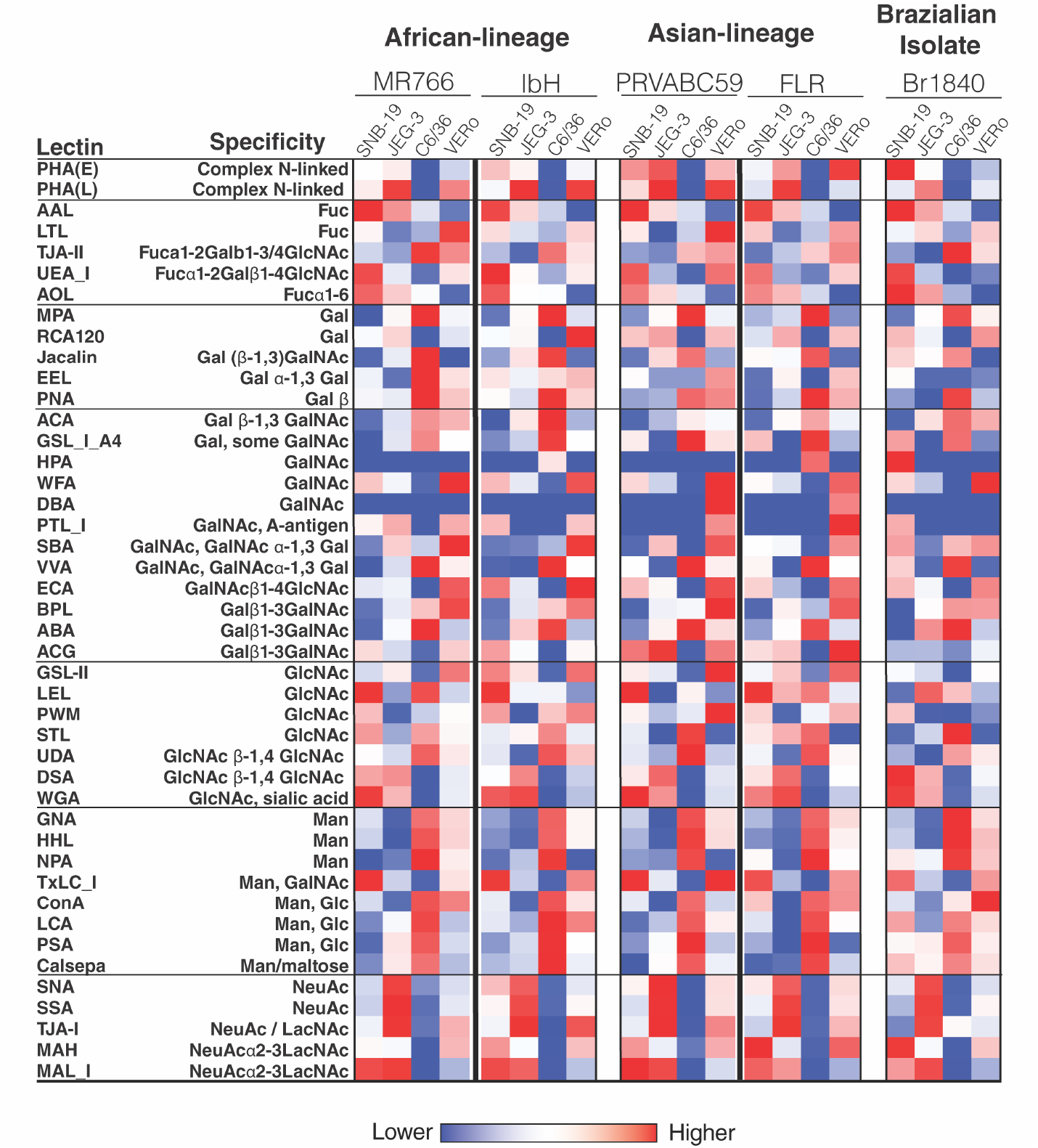
Glycomic profiling of ZIKV-infected and uninfected cell membrane using a lectin microarray. A lectin microarray (45 structures) was used to analyze the glycans associated to cell-surface proteins in both ZIKV-infected and uninfected cells. The different cell types (SNB-19, JEG-3, C6/36 and Vero cells) were either mock-infected or infected with distantly related ZIKV strains (MR766, IbH, PRVABC59, FLR, and SJRP). The cell-surface proteins were later extracted using the membrane protein extraction kit, labeled with Cy3 dye and hybridized to a lectin microarray. The resulting lectin chip was scanned for fluorescence intensity on each lectin-coated spot using an evanescent-field fluorescence scanner. The data were normalized using the global normalization method. The heatmap format was used to present the glycan profiles identified using this method. Man represents high-mannose, GalNAc represents N-acetylgalactosamine, GlcNAc represents N-acetylglucosamine, NeuAc represents N-acetylneuraminic acid, Gal represents galactose, Glc represents glucose, and Fuc represents fucose.

**Fig. 4.**
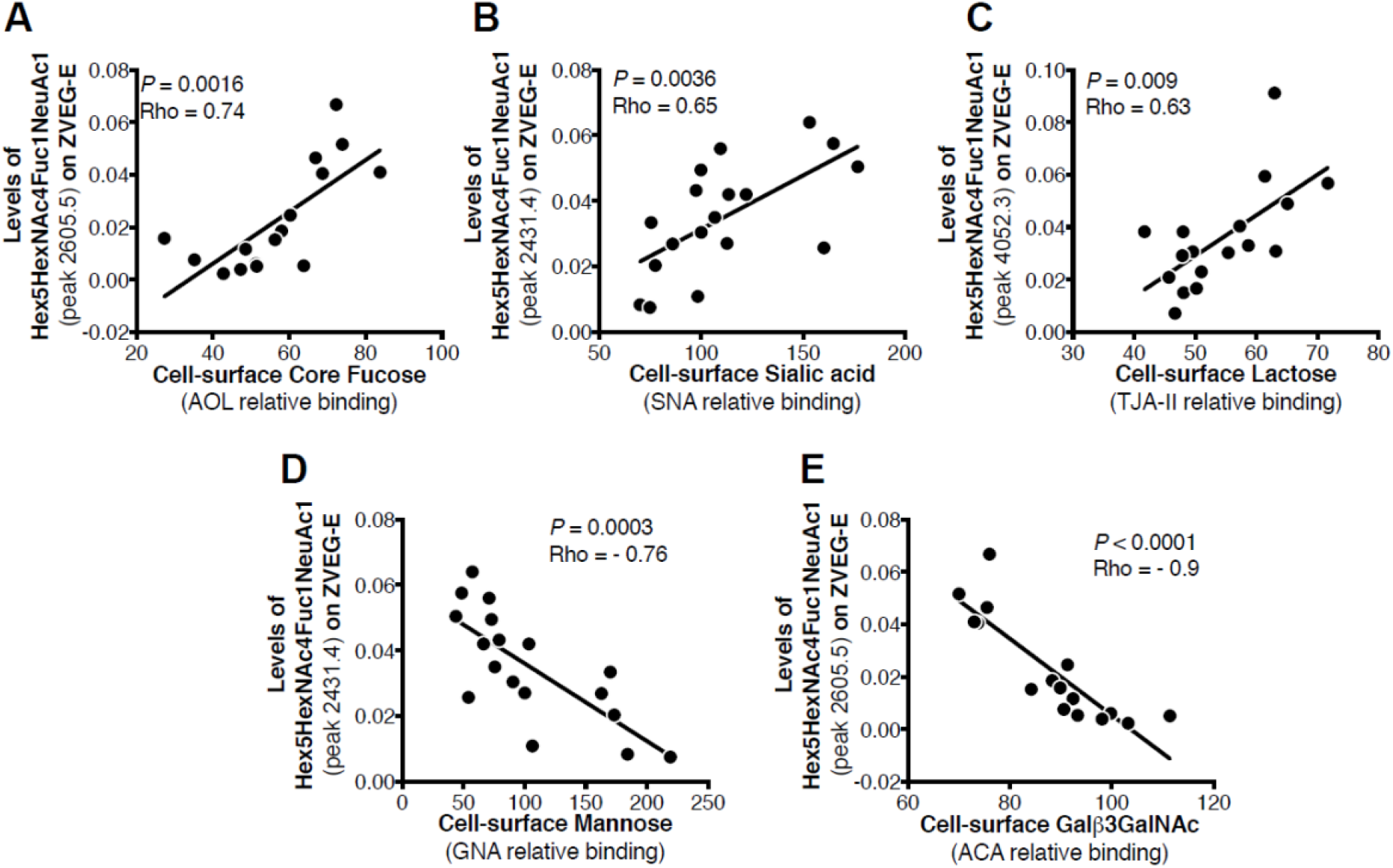
Comparison of glycan profiles identified by MALDI-TOF-MS and by lectin microarray. A significant linear correlation was confirmed between the cell-surface glycans identified by the lectin microarray (fucose, sialic acid, lactose, mannose, and GlcNAc) and the glycomic features of Zika virus protein E that were observed through MALDITOF-MS.

### C-type Lectins DC-SIGN and L-SIGN play a functional important role in ZIKV infection

The C-type lectins such as DC-SIGN (dendritic cell-specific ICAM-grabbing non-integrin, where ICAM is an intercellular adhesion molecule) and L-SIGN (where L is liver or lymph node) have considerable binding affinity to viral mannose-rich glycan ^37,38^. DC-SIGN is expressed on various cell types such as monocytes and dendritic cells (DCs), and L-SIGN is expressed on endothelial cells of liver and lymph nodes (L) and recognizes carbohydrate structures present on viral glycoproteins ^38^. Thus, these lectins function as attachment factors for several enveloped viruses including human immunodeficiency virus-1 (HIV-1), Ebola (EBOV), Japanese encephalitis virus (JEV), and Dengue virus (DENV) ^39,40^. Given that ZIKV envelope-associated N-glycan structures are enriched with mannose, we examined the potential interactions of ZIKV E glycans with host lectins (DC-SIGN and L-SIGN), and their impact on ZIKV entry and infection. We employed 3T3/NIH cells (which were originally established from the mouse embryonic fibroblast cells) expressing DC-SIGN (DC-SIGN/3T3 cells) and L-SIGN (L-SIGN/3T3 cells) receptors, along with only 3T3/NIH cells as a control. Cells were infected with ZIKV at a MOI of 1, and the percent infection was determined after 24 hours through the use of an immunofluorescence analysis and a high-throughput Operetta imager. As presented in **Fig. 5**, the percentage (%) of infected cells increased in DC-SIGN and L-SIGN expressing cells compared to control 3T3/NIH cells, indicating that DC- and L-SIGN expressing 3T3 cells are susceptible to ZIKV infection **(Fig. 5a)**. To further confirm the role of high-mannose type glycans, we performed experiments in the presence of mannan and EDTA. EDTA blocks DC-SIGN by extracting the bound calcium, while Mannan competes with insect-derived high-mannose type glycans and both inactivate DC-SIGN ^41^. Both mannan and EDTA inhibited ZIKV replication efficiently **(Fig. 5b, c)**. These results suggest that ZIKV infection efficiency is dependent upon high-mannose type glycans, and the entry is mediated through DC-SIGN/L-SIGN cell surface expression. Taken together, DC-SIGN/L-SIGN receptors are essential for optimizing the entry and infection of various strains of ZIKV.

**Fig. 5.**
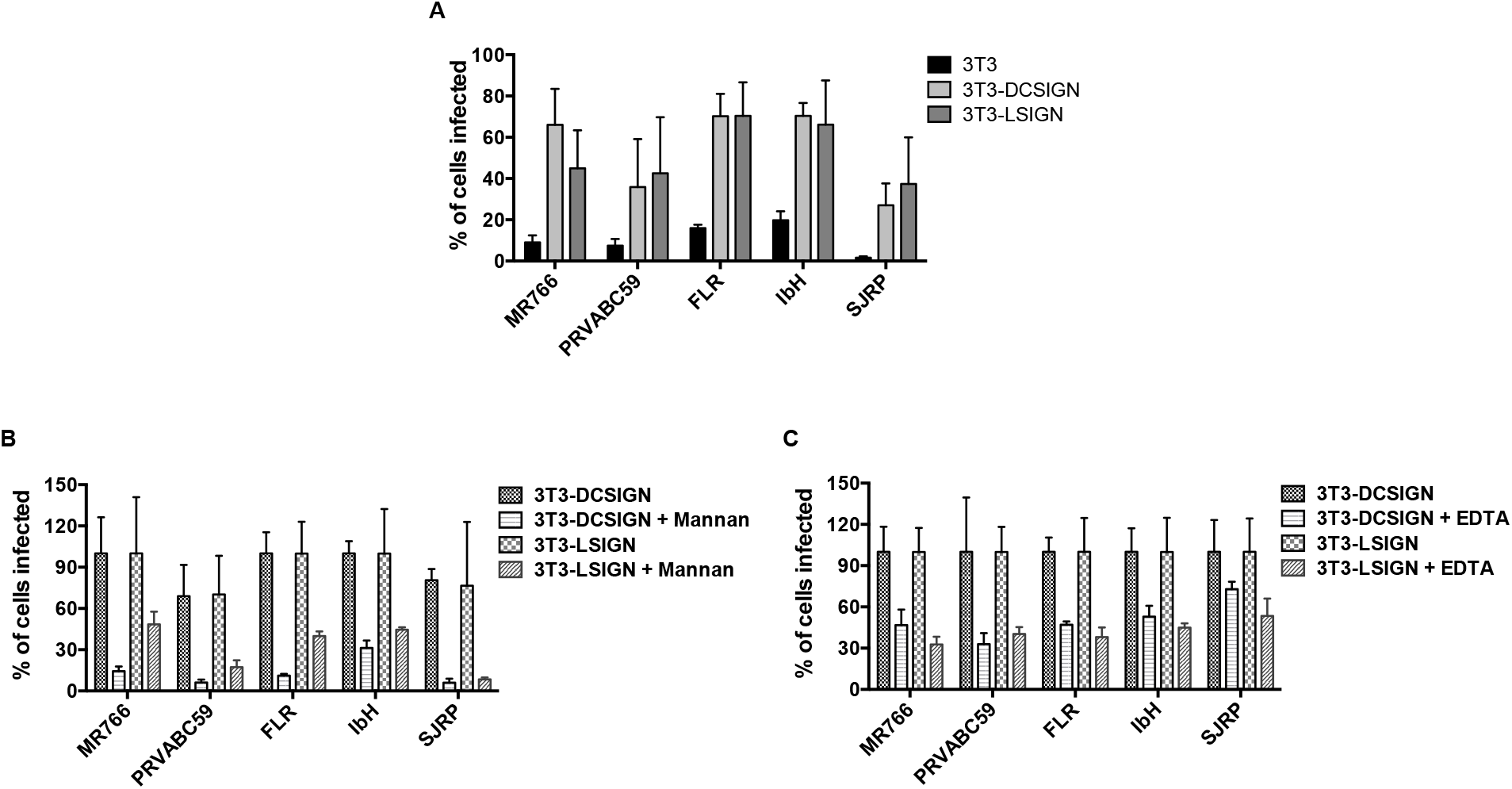
The functional importance of N-linked glycans on Zika virion infectivity. (a) DC-SIGN/L-SIGN-mediated enhancement of Zika virus infection. 3T3, 3T3-DC-SIGN and 3T3-L-SIGN cells were infected with ZIKV at MOI of 1, and the expression of viral E glycoprotein was detected 24 hours later through the use of an immunofluorescence assay. (b) DC-SIGN/L-SIGN-mediated enhancement of Zika virus infection is inhibited by Mannan. 3T3-DC-SIGN or 3T3-L-SIGN cell lines were incubated with Mannan (at 200 μg/ml) inhibitors for 30 min at 37 °C and then infected with ZIKV at MOI of 1. The same concentration was maintained constant throughout duration of infection. (c) DC-SIGN/L-SIGN-mediated Zika virus infection is reduced by EDTA. 3T3-DC-SIGN or 3T3-L-SIGN cell lines were preincubated with EDTA (at 0.5 mM) for 30 min at 37 °C and then infected with ZIKV at MOI of 1. The treatment was continued for 2 h. After 2h, the virus and the EDTA containing medium was removed and continued the infection with the fresh medium. After 24h, the infected cells were detected using viral E protein antibody and the percent of infected cells was assessed by an immunofluorescence assay using Operetta High-Content Imaging System. Error bars indicate standard deviations calculated from 4 replicate wells in a 96-well plate.

## Discussion

A comprehensive understanding of glycosylation pattern of the viral antigenic proteins is a key towards therapeutic discovery and vaccine development. It is equally important to understand how glycosylation affects the biological properties and pathogenesis of the virus, and for designing glycomimetic compounds as potential antiviral agents ^42^. In this study, we used MS to characterize the E protein N-glycans of ZIKV derived from insect cells and mammalian cells. We showed for the first time that ZIKV E, which is required for host cell receptor binding, is differentially glycosylated in a cell-type different fashion. The N-glycosylation pattern on ZIKV E is thought to be involved in viral infectivity, morphogenesis, protein folding, virus entry, and tissue tropism among other functions. This has also been described for other viruses including influenza virus ^43^, HIV-1, West Nile Virus, JEV, hepatitis C virus (HCV), Nipah virus, EBOV, and sever acute respiratory syndrome (SARS) coronavirus ^44^ Analyses of the N-glycans on ZIKVs grown in insect and various mammalian cells revealed a high degree of heterogeneity. The variety of N-glycans found on ZIKV E ranged from high- (or oligo-) mannose, notably Man5GlcNAc2, to highly sialylated (up to four sialic acids) complex-type N-glycans and with very few hybrid types. However, both African and Asian lineage strains revealed very similar N-glycosylation patterns of ZIKV E. Di- and tri-sialylated bi- and tri-antennary were the most relatively abundant N-glycans found on ZIKV E when viruses were produced in C6/36 insect cells and LLC-MK2, Vero, THP-1, SNB19, and JEG-3 mammalian cells. High-mannose, asialo- and agalacto-N-glycans were also identified, though in lesser overall relative abundance. The presence of high-mannose and complex-type N-glycans on mammalian cell-derived virus is not limited to ZIKV as has been reported for DENV serotypes ^45,46^, and is the most abundant type of N-glycans found on HIV GP120 glycoprotein^47^. On the other hand, any ZIKV strain when produced in various cell lines showed different N-glycan patterns. Indeed, from a cell line perspective, LLC-MK2 cells produced the same unique N-glycosylation pattern for all five ZIKV E, which were mostly decorated with bi-antennary di-sialylated and core-fucosylated agalactosylated N-glycans **(Fig. 2-3, Supplementary Fig. 4**). Collectively, these findings indicate that not only virus characteristics; *e.g*. strain or lineage, but also host cell physiology has an impact on the glycosylation pattern of ZIKV E which may affect viral pathogenesis of ZIKV strains.

In the current study, we did not determine the location of the N-glycans on ZIKV E protein. We predict that the processed complex (mammalian-derived virus) or paucimannose (insect-derived virus) glycan is at position N154 which is not essential for virus production and spread in mosquito or mammalian cells ^48^, as the high mannose-type N-glycans have been reported to be present at the N154 residue in DI domain of ZIKV E within the conserved glycosylation motif N-X-S/T ^25,49^. Moreover, Asian strains (including strains circulating in Southeast Asia, and South and Central America) of ZIKV, but not African strains, contain the N154 glycosylation site of the E protein (**Supplementary Fig. 1**), which has been reported previously ^50^. Contrasting the four African and Asian lineage viruses, the Brazilian ZIKV isolate SJRP revealed only partial similarities in N-glycosylation. SJRP was decorated only with non-sialylated N-glycans when generated in both C6/36 and Vero cell lines, however, mostly with sialylated N-glycans in THP-1, SNB-19 and JEG-3 cells **(Fig. 2-3, Supplementary Fig. 4**), suggesting that decoration of ZIKV E with sialylated N-glycans may correlate with diverse tissue tropism of the virus. Taken together, these findings lead us to hypothesize that the “N-glycotype” of the recently emergent Asian strains is mediating the incidence of the new neuropathogenic potential of ZIKV infection.

The results of our study revealed high heterogeneity in structure and composition of the N-glycans linked to ZIKV E produced by both insect and various mammalian cells including monocytes, placental, and neural cell types. The heterogeneity of N-glycan has also been linked to DENV E protein in terms of structure and composition ^23^. The N-glycan on DENV E protein that is produced by mammalian cells is a mixture of high-mannose glycan and complex glycan. Mosquito cells, by contrast, provide a mix of high-mannose glycan and paucimannose glycan ^46^. It has been shown that ZIKV replication in macaques also restores the absent N-glycosylation site in MR766 strain ^51^, suggesting that the passage history (host animal and cell line types) affects E protein glycosylation pattern of ZIKV strains. Taken together, our study revealed glycan structures linked to ZIKV E, analyzed by using MS and lectin microarray, are varied in structure and composition depending on ZIKV strains and host cell types.

The C6/36, Vero, THP-1, and SNB-19 cells produced ZIKV E N-glycosylation profiles with highly sialylated N-glycans on the E protein of all ZIKV strains, except for MR766 in JEG-3 cell and SJRP in C6/36 and Vero cells where the N-glycosylation profile was mostly restricted to low molecular weight non-sialylated N-glycans. These data suggest that the different contexts of specific host cells could significantly affect the N-glycosylation profile of the ZIKV E of different ZIKV strains. The cell culture media, culture conditions, and sequence of a protein may also significantly influence the quality and relative abundance of N-glycans observed on the cell-produced glycoproteins. The results of our study also revealed a high heterogeneity in the N-glycans linked to ZIKV E among different ZIKV strains that are produced by the same host cell. For example, Vero cells produced ZIKV E linked to various N-glycans among ZIKV strains including FRL, IbH, PRVABC59, and MR766. Unlike other flaviviruses, ZIKV E contains a conserved sequence of ~ ten amino acids that surround the N154 glycosylation site. This conserved sequence may affect the glycosylation process, carbohydrate moiety, and attachment site of the ZIKV to host cells ^18^. The number and location of glycosylation motifs in the E vary considerably both between and within flaviviruses ^52^, altering the pathogenicity of these viruses and the severity of their diseases. The African lineage of ZIKV isolates lack the N-glycosylation site while ZIKV isolates from recent outbreaks in South and Central American regions, that was originated from the epidemy of Asian lineage ZIKV PF-2013 in French Polynesia in 2013, contain the glycosylation site within the viral envelope ^22^. Indeed, in a previous study, the recombinant ZIKV strains that lack the glycosylation site were unable to induce mortality in the mouse model ^22^. However, recent studies revealed that residues surrounding the ZIKV E protein glycan regulate virus antigenicity, irrespective of the presence of a glycan ^53^. The glycosylation of ZIKV E protein does not affect antibody binding to a nearby epitope or its capacity to serve as a neutralization target ^54^ Collectively, these findings suggest that ZIKV pathogenicity, but not immunogenicity, might be influenced by the glycosylation of the E protein which occurs post-translationally in ER of the host cell. This host cell-dependent glycosylation process of ZIKV E could thus have wide ranging implications regarding the pathogenicity and infectivity of the virus. However, future research is required to evaluate the full extent of this circumstance.

In regard to infection of cells, we explored ZIKV interactions with cellular lectins that bind specific types of glycans we identified on the virus. DC-SIGN is a lectin-like molecule, expressed in dendritic cells that can facilitate virus entry through its binding affinity with the glycans on the E protein ^55^. N154 glycosylation was found to play a role in ZIKV infection of mammalian cells through this entry factor DC-SIGN ^48^. Indeed, the glycosaminoglycan (GAGs) of C-type lectins, including DC-SIGN, have a binding affinity as an attachment factor for host cell entry of ZIKV similar to other pathogenic flaviviruses ^56^. We also observed a significant decrease in ZIKV infection in DC-SIGN and L-SIGN expressing cells in the presence of competitive inhibitors rich in mannose. As a result, the presence of monoclonal antibodies to various C-type lectins, the competitive inhibitors rich in mannose, or antiviral gene therapy may be able to reduce ZIKV replication by impairing the binding between viral N-glycan of E protein and GAGs of DC-SIGN. Recent reports suggest that the cells expressing DC-SIGN, such as human dermal fibroblasts, epidermal keratinocytes, and immature dendritic cells, are permissive to the most recent ZIKV isolates. This may have contributed to the epidemic in French Polynesia in 2013 where the cell entry mechanism of ZIKV strain PF-2013 to various cell lines was shown to be mediated by a range of receptors and molecules including DC-SIGN ^57^ The accessibility of ZIKV PF-2013 strain, the original strain initiated in the recent outbreaks of ZIKV in South and Central American regions, to different entry cell receptors possibly provides an evolutionary advantage for this recent ZIKV strain to infect a wide range of target cells and evade the immune system of human host. Taken together, these phenomena may partially explain the alterations in the mechanisms of pathogenesis and tissue tropism of these recent ZIKV strains. Further studies are needed to determine the impact of host glycosylation variability on ZIKV pathogenesis and infectivity *in vivo*.

In conclusion, our findings provide a first-of-its-kind detailed repertoire of N-glycans linked to the ZIKV envelope. To our knowledge, this is the first comprehensive mapping on the glycome of ZIKV E in different physiologically relevant cell lines. Our study showed that the N-glycans linked to ZIKV E produced by both insect and various mammalian cells are highly heterogenous in structure and composition. Both African and Asian lineage strains showed very similar N-glycosylation patterns of ZIKV E. We also noticed that the N-glycan pattern of the ZIKV E depends on not only virus strain characteristics but also on cell line type. We showed that ZIKV infection in DC-SIGN and L-SIGN expressing cells was significantly decreased in the presence of competitive inhibitors rich in mannose, suggesting an important role for these lectin-like molecules as cell entry factors in pathogenesis, emerging new strains of ZIKV and alternative targets for therapeutic development. The results of this study are important in efforts to design therapeutically active antibodies against ZIKV replication and glycomimetic compound-based antivirals.

## Methods

### Cell lines and viruses

The ZIKV stocks were produced in various cell lines derived from human (placenta, brain, and monocytes), monkey, and insects **(Supplementary Table 1a)**, and tittered in Vero (kidney epithelial cells extracted from an African green monkey) cells. ZIKV strains, MR766 (Rhesus/1947/Uganda; BEI Cat. # NR-50065),PRVABC59 (Human/2015/Puerto Rico; BEI Cat. # NR-50240), FLR (Human/2015/Colombia; BEI Cat. # NR-50183), IbH 30656 (Human/1968/Nigeria; BEI Cat. # NR-50066) were obtained from BEI Zika resources. A ZIKV Brazilian isolate (SJRP-HB-2016-1840 – herein referred to as SJRP) was also obtained from University of Texas Medical Branch (UTMB) Arbovirus reference collection **(Supplementary Table 1b)**. The cells were cultured and maintained in Dulbecco’s modified eagle medium (DMEM, Gibco, Life Technologies, Carlsbad CA, USA) supplemented with 10% heat-inactivated fetal bovine serum (FBS, Invitrogen, Carlsbad, CA, USA) at 37 °C in a 5% carbon dioxide humidified environment. The mammalian cell lines JEG-3 (human placental choriocarcinoma) and SNB-19 (human brain glioblastoma) were grown on DMEM (1X) supplemented with the 10% FBS. The LLC-MK2 (kidney epithelial cells extracted from *Macaca mulatta* monkey) cells were grown on Medium 199 (Biowest, Riverside, MO, USA) supplemented with 1% horse serum (Invitrogen, CA, USA). The THP-1 (human monocytic cells) cells were grown on Roswell Park Memorial Institute (RPMI) medium (Gibco, CA, USA) supplemented with 10% FBS and 0.05 mM β-mercaptoethanol (Sigma-Aldrich, St. Louis, MO, USA). The mosquito C6/36 (*Aedes albopictus* clone) cells were grown on minimum essential medium (MEM, Gibco, Life Technologies, CA, USA) supplemented with 10% FBS. All cell culture media were supplemented with 1% penicillin-streptomycin (Gibco, CA, USA) and maintained at 37 °C in a 5% carbon dioxide humidified environment, except the C6/36 cells which were maintained at 28 °C.

### Virus infection

The various ZIKV strains, such as African strain (MR766 and IbH), Asian strains (PRVABC59 and FLR), and a primary isolate from Brazil in 2016 (SJRP) at multiplicity of infection (MOI) of 1 was adsorbed onto cells. After 90 min, in serum-free medium with rocking every 15 min, the inoculum was removed; the C6/36, Vero, LLC-MK2, SNB-19, THP-1, and JEG-3 cells were maintained in respective medium containing 2% FBS containing respective medium. All the cells lines were maintained at 37 °C in a CO2 incubator (except the C6/36 cells which were maintained at 28 °C). For the large-scale virus production, all cells were maintained at 37 °C in CO2 incubator for 3-4 days (except C6/36 cells which were maintained at 28 °C for 5-7 days) until cytopathic effects were observed.

### Immunofluorescence assay

The C6/36, Vero, and SNB-19 cells were grown on cover slips and mock-infected or infected with various strains of ZIKV. After 24 hours post infection, cells were processed for indirect immunofluorescence assay using the double-labelling method. Briefly, cells were fixed in 4% paraformaldehyde and processed for immunofluorescence assay. The cells were blocked and permeabilized using 5% goat serum (Sigma-Aldrich, St. Louis, MO, USA) with 0.5% Triton X-100 (Sigma-Aldrich, St. Louis, MO, USA). The cells were then immunoassayed using rabbit polyclonal antibody (1:1000) against the E protein of ZIKV (Genetex Inc., Irvine, CA, USA), in a combination with ER, using ER ID dye as per manufacturer’s instructions (Enzo Life Sciences, Inc, Farmingdale, NY, USA) and the cell nuclei (1:2000), using Hoechst stain (Invitrogen, Rockford, lL, USA) for 15 min. Subsequently, Alexa Fluor 594-conjugated secondary antibody (1:2000) (Invitrogen, Rockford, IL, USA) was added onto cells; and images were captured under 40X using Operetta High-Content Imaging System (PerkinElmer, Waltham, MA, USA).

### Purification of ZIKV virus and isolation of E protein

A large stock of ZIKV viruses were purified using sucrose cushion in ultracentrifugation as described in previous methods ^26^. Briefly, culture supernatants were centrifuged at 6,000 rpm for 10 min, and a 0.45 μm filter was used to remove cell debris. Virus-containing filtrate was layered onto 20% (w/v) sucrose and subjected to ultracentrifugation at 100,715 × g, 4 °C for 3.5 hours. Pellets were dissolved in 100 μl of NTE buffer (10 mM Tris-HCl, pH 8.0, 120 mM NaCl and 1 mM EDTA (ethylene diamine tetra acetic acid)). The purified virions were then resolved on gradient 4-12 % bis-tris protein gels. The presence of ZIKV glycoprotein E was confirmed by western blot analysis, and confirmed band was excised for MS analysis.

### Lectin microarray

For the lectin array analysis, the cells (C6/36 cells, Vero cells, SNB-19 cells, and JEG-3 cells) were grown to 60-70 % confluency. Various ZIKV strains, such as African strain (MR766 and IbH), Asian strains (PRVABC59 and FLR) and the primary Brazilian isolate SJRP, at MOI of 1 were then added onto cells for 90 min, in serum-free medium. After 90 min, the inoculum was removed; and cells continued to culture in their respective culture conditions. The cells were harvested using cell scrapper, made into a single cell suspension, and counted using cell countess (Invitrogen, Carlsbad, CA, USA). The lectin microarray enabled sensitive analysis of multiple glycan structures (45 structures) by employing a panel of immobilized lectins with known glycan binding specificity ^27–35^. Cell-surface proteins were purified using Mem-PER™ Plus Membrane Protein Extraction Kit (Thermo Fisher Scientific, Rockford, lL, USA). Isolated proteins were labeled with Cy3 dye and hybridized to a lectin microarray ^27–35^. The resulting lectin chips were scanned for fluorescence intensity on each lectin-coated spot using an evanescent-field fluorescence scanner. Data were normalized using the global normalization method.

### N-glycan preparation

Gel bands stained with Coomassie Blue were excised and transferred into clean tubes. Then, 200 μl of 50 mM ammonium bicarbonate (AMBIC) (Sigma-Aldrich, St Louis, MO, USA) and 200 μl of acetonitrile (Sigma-Aldrich, St. Louis, MO, USA) were added to the samples, mixed thoroughly and incubated at room temperature (RT) for 5 min. The supernatants were then discarded, and the washing step was repeated once more. The gel pieces were dried with a vacuum centrifuge for 10 min. Then 200 μl of a 10 mM DTT (1,4-Dithiothreitol, Sigma-Aldrich, St. Louis, MO, USA) solution was added and incubated at 50 °C for 30 min. The DTT solution was discarded, and the samples were briefly washed with 200 μl of acetonitrile solution. Again, the samples were dried with a vacuum centrifuge for 10 min.

After drying, the samples were incubated with 200 μl of a 55 mM IAA (Iodoacetamide, Sigma-Aldrich, St. Louis, MO, USA) solution for 30 min in the dark at RT. The IAA solution was then discarded, and the samples were washed with 500 μl of 50 mM AMBIC for 15 min at RT, followed by 5 min incubation with 200 μl of acetonitrile. The samples were dried once again with a vacuum centrifuge for 10 min, before adding 500 μl of 50 mM AMBIC containing 10 μg of TPCK-treated trypsin (Sigma-Aldrich, St. Louis, MO, USA). After incubating overnight at 37 °C, the trypsin digestion was terminated by boiling the sample for 3 min.

The supernatants were recovered and collected in a clean glass tube, before carrying out two sequential washes with 200 μl of 50 mM AMBIC, vortexed for 15 min; 200 μl of 50% acetonitrile in 50 mM AMBIC, vortexed for 15 min; and 200 μl of acetonitrile, vortexed for 15 min. For each sample, all washes were collected, pooled in the same glass tube that was previously used, and then lyophilized. The dried materials were resuspended in 200 μl of 50 mM AMBIC. 1 μl of PNGaseF (New England Biolabs, Ipswich, MA, USA) was added to this for an overnight incubation at 37 °C. Two drops of 5% acetic acid (Fisherbrand, Waltham, MA, USA) were added to stop the enzymatic reaction before purifying the released N-glycan over a C18 Sep-Pak (50 mg) column (Waters, Milford, MA, USA) that was conditioned with 1 column volume (CV) of methanol (Sigma-Aldrich, St. Louis, MO, USA), 1 CV of 5% acetic acid, 1 CV of 1-propanol (Sigma-Aldrich), and 1 CV of 5% acetic acid. The C18 column was washed with 3 ml of 5% acetic acid, flow through; and wash fractions were collected, pooled, and lyophilized.

### Permethylation of N-glycan

Lyophilized N-glycan samples were incubated with 1 ml of a DMSO (Dimethyl Sulfoxide; Sigma)-NaOH (Sigma-Aldrich, St. Louis, MO, USA) slurry solution and 500 μl of methyl iodide (Sigma-Aldrich, St. Louis, MO, USA) for 20-30 min under vigorous shaking at RT. 1 ml of Milli-Q water was added to stop the reaction. 1 ml of chloroform (Sigma-Aldrich, St. Louis, MO, USA) was added to purify the permethylated N-glycan. 3 ml of Milli-Q water were added to wash the chloroform fractions, and the mixture was briefly vortexed. The water was discarded by additional centrifugation. This wash step was repeated 3 times. The chloroform fraction was dried before being redissolved in 200 ml of 50% methanol. This was then loaded into a conditioned (1 CV methanol, 1 CV Milli-Q water, 1 CV acetonitrile (Sigma-Aldrich, St. Louis, MO, USA), and 1 CV Milli-Q Water) C18 Sep-Pak (200 mg) column. The C18 column was washed with 6 ml of 15% acetonitrile and then eluted with 6 ml of 50% acetonitrile. The eluted fraction was lyophilized and then redissolved in 10 μl of 75%methanol from which 1 μl was mixed with 1 μl DHB (2,5-dihydroxybenzoic acid (Sigma-Aldrich, St. Louis, MO, USA)) (5mg/ml in 50% acetonitrile with 0.1% trifluoroacetic acid (Sigma-Aldrich, St. Louis, MO, USA)) and spotted on a MALDI polished steel target plate (Bruker Daltonics, Bremen, Germany).

### MS data acquisition and analyses

MS data were acquired on a Bruker UltraFlex II MALDI-TOF Mass Spectrometer instrument. The reflective positive mode was used, and data were recorded between 500 m/z and 6000 m/z. For each MS N-glycan profile, the aggregation of 20,000 laser shots or more were considered for data extraction. Mass signals of a signal/noise ratio of at least 4 were considered, and only MS signals matching an N-glycan composition were considered for further analysis. Subsequent MS post-data acquisition analysis was made using mMass^36^. The relative abundance of each N-glycans identified on ZIKV E in each experimental condition was calculated based on the absolute intensity of the first isotopic peak of a given N-glycan relative to the sum of all N-glycan intensities.

### Assays to evaluate the functional importance of N-linked glycans

In inhibition experiments, 3T3, 3T3-DCSIGN and 3T3-L-SIGN cells were incubated with 200 μg/mL of yeast mannan (Sigma-Aldrich, St. Louis, MO, USA) or 0.5 mM EDTA at 37 °C for 30 min. Various strains of Zika viruses (MOI = 1) were preincubated with same concentration of inhibitor and adsorbed on pretreated cells. The mannan treatment was maintained constant throughout duration of infection. Whereas, the EDTA and virus containing medium was removed after 2h and continued the infection with the fresh medium. The infected cells were detected using viral E protein antibody, and the percent of infected cells was assessed by an immunofluorescence assay using Operetta High-Content Imaging System (PerkinElmer, Inc., Waltham, MA, USA).

### Statistical Analysis

Spearman’s Rank Order Correlations were conducted using GraphPad Prism release 7.0 (GraphPad Software, San Diego, CA, USA).

## Acknowledgments

We thank Russell Jaffe and Robin Taylor for editorial assistance. We acknowledge Krishna Kota (USAMRIID) for his help with Operetta high-content imaging system. We thank Dr. Nikos Vasilakis (UTMB) for kindly providing Zika Brazilian isolate SJRP-HB-2016-1840. This work is supported in part by R01AI113883, Nebraska Neuroscience Alliance Endowed Fund Award to SNB, and the National Center for Functional Glycomics Grant P41GM103694 to RDC.

## Author information Affiliations

Department of Pharmacology and Experimental Neuroscience, University of Nebraska Medical Center, Omaha, NE 68198-5800, USA

Nanda Kishore Routhu, Mehdi R. M. Bidokhti & Siddappa N. Byrareddy

Department of Genetics, Cell Biology and Anatomy, University of Nebraska Medical Center, Omaha, NE 68198-5805, USA; Department of Biochemistry and Molecular Biology, University of Nebraska Medical Center, Omaha, NE 68198-5805, USA

Siddappa N. Byrareddy

Beth Israel Deaconess Medical Center, National Center for Functional Glycomics, Boston, MA, USA

Sylvain D. Lehoux & Richard D. Cummings

Beth Israel Deaconess Medical Center Glycomics Core, Boston, MA, USA; 6The Wistar Institute, Philadelphia, PA, USA

Sylvain D. Lehoux & Emily A. Rouse

Department of Pathology and Microbiology, University of Nebraska Medical Center, Omaha, NE, USA.

St Patrick Reid

The Wistar Institute, Philadelphia, PA, USA

Leila B. Giron, Alit Anzurez & Mohamed Abdel-Mohsen

## Author Contributions

N.K.R. designed, performed, analyzed the data, and drafted the manuscript. S.N.B. designed and supervised the study, contributed to data analysis, and edited the manuscript. S.D.L., and E.A.R. performed the MS analyses. S.D.L., E.A.R., and R.D.C. analyzed the MS data. M.A.M., and L.B.G., performed the lectin array. M.R.M.B., and P.C.R. helped in manuscript editing and interpretation. All authors provided critical feedback and helped shape the research, analysis and manuscript editing.

## Corresponding author

Correspondence to Siddappa N. Byrareddy.

## Supplementary Information (Word file)

